# AFP2 inhibits ABA responses during germination without ABI5 degradation but DWAs reduce desiccation tolerance

**DOI:** 10.1101/2021.12.21.473709

**Authors:** Tim Lynch, Guillaume Née, Avan Chu, Thorben Krüger, Iris Finkemeier, Ruth R Finkelstein

## Abstract

Overexpression of ABI5/ABF interacting proteins (AFPs) results in extreme ABA resistance of seeds and failure to acquire desiccation tolerance, at least in part through effects on chromatin modification. This study tests the hypothesis that the AFPs promote germination by also functioning as adapters for E3 ligases that ubiquitinate ABI5, leading to its degradation. Interactions between AFPs and two well-characterized classes of E3 ligases targeting ABI5, <u>DW</u>D HYPERSENSITIVE TO <u>A</u>BA (DWA)s and KEEP ON GOING (KEG), were analyzed by yeast two-hybrid, bimolecular fluorescence complementation, and genetic assays. Although the AFPs and E3 ligases showed weak direct interactions, loss of function for the E3 ligases did not impair ABA-resistance conferred by overexpression of the YFP-AFP2 fusion. Comparison of ABI5 and AFP2 levels in these lines showed that AFP2 accumulation increased during germination, but that ABI5 degradation followed germination, demonstrating that AFP2 controls ABA sensitivity during germination independently of ABI5 degradation. Surprisingly, AFP2 overexpression in the *dwa1 dwa2* mutant background produced the unusual combination of extreme ABA resistance and desiccation tolerance, creating an opportunity to separate the underlying biochemical characteristics of ABA sensitivity and desiccation tolerance that we investigated by quantitative proteomics. Our analysis identified at least three-fold more differentially accumulated seed proteins than previous studies. Comparison of dry seed proteomes of the different genotypes allowed us to separate and refine the changes in protein accumulation patterns correlating with desiccation tolerance independently of ABA sensitivity, or *vice versa*, to a subset of cold-induced and defense stress-responsive proteins and signaling regulators.

**Summary Sentence:** Extreme ABA resistance conferred by overexpression of AFP2 is not mediated by interactions with E3 ligases, but the *dwa* background maintains desiccation tolerance despite ABA resistance.

## Introduction

Germination inhibition is a well-characterized response to ABA that has been used in many screens for mutants with defects in ABA response. The ABA-INSENSITIVE(ABI)5 locus was identified through such a screen (Finkelstein, 1994), and subsequently found to encode a bZIP transcription factor that regulates gene expression in maturing seeds and seedlings (Finkelstein and Lynch, 2000) and a “developmental checkpoint” controlling the transition from germination to seedling growth (Lopez-Molina et al., 2001). Numerous studies have demonstrated a correlation between ABI5 accumulation in response to stress following imbibition and failure to pass this checkpoint (Stone et al., 2006; Piskurewicz et al., 2008; Lee et al., 2010; Albertos et al., 2015; Xu et al., 2019; Ji et al., 2019). However, these studies did not distinguish between ABI5 accumulation as cause or effect of non-germination.

Early efforts to understand factors regulating ABI5 accumulation and activity led to identification of proteins including related and distinct classes of transcription factors, kinases and phosphatases, E3 ligases (reviewed in (Cutler et al., 2010; Finkelstein, 2013; Yu et al., 2015; Skubacz et al., 2016) and a highly conserved family of ABI5-interacting proteins (AFPs) (Lopez-Molina et al., 2003; Garcia et al., 2008) whose homologies were initially characterized only as domains of unknown function (DUFs). Many possible functions have been proposed for the AFPs, including acting as transcriptional co-repressors affecting chromatin structure (Pauwels et al., 2010; Lynch et al., 2017) or, in the case of AFP1, adapters for E3 ligases that add ubiquitin to ABI5 (Lopez-Molina et al., 2003). These roles are not mutually exclusive and recent analyses have suggested that the AFPs may be intrinsically unstructured proteins (Lynch et al., 2017), consistent with the ability to assume a variety of conformations allowing them to interact with diverse proteins, possibly acting as scaffolds.

Two well-characterized classes of E3 ligases that directly negatively regulate ABI5 accumulation are the <u>DW</u>D HYPERSENSITIVE TO<u>A</u>BA (DWA) proteins (Lee et al., 2010) and <u>KE</u>EP ON <u>G</u>OING (KEG)(Stone et al., 2006). DWAs are redundantly acting nuclear proteins that ubiquitinate ABI5 such that *dwa1dwa2* mutants are hypersensitive to ABA for inhibition of germination and this is correlated with increased ABI5 accumulation (Lee et al., 2010). Loss of KEG has a similar effect on ABA sensitivity and ABI5 accumulation, but KEG is associated with the trans-Golgi network. Consequently, KEG interactions with ABI5 are restricted to the cytoplasm (Liu and Stone, 2013), and are promoted by S-nitrosylation of ABI5 (Albertos et al., 2015). KEG is a RING-type E3 ligase that auto-ubiquitinates in the presence of ABA, leading to its own destruction via the proteasome, thereby permitting stabilization of ABI5 that can be transported into nuclei (Liu and Stone, 2010). More recently, stabilization of ABI5 within nuclei was found to depend on association with the XPO1-interacting WD40 protein (XIW)1 (Xu et al., 2019). Although the initial suggestion that AFPs acted as adapters for E3 ligases was based on colocalization of AFP1 and CONSTITUTIVELY PHOTOMORPHOGENIC (COP)1 (Lopez-Molina et al., 2003), COP1 was recently shown to positively regulate ABA inhibition of seedling establishment by promoting ABI5 binding to its target promoters (Yadukrishnan et al., 2020), possibly indirectly through either COP1-mediated destruction of proteins that antagonize ABI5 function (Xu et al., 2016) or a COP1-HY5-ABI5 regulatory module (Chen et al., 2008). However, no significant changes in ABI5 transcript or protein accumulation were detected in *cop1* mutant seedlings.

In this study, we tested whether ABI5 degradation is required for AFPs to promote ABA-resistant germination. To determine whether AFPs serve as adapters for either the DWA or KEG class of E3 ligases, we tested for epistatic interactions between mutations in these E3 ligases and AFP2 overexpression, which had previously been shown to confer an approximately 100-fold increase in resistance to ABA and a resulting failure to complete seed maturation (Lynch et al., 2017). All mutant combinations remained resistant to ABA despite maintaining ABI5 accumulation until germination was nearly complete, showing that ABI5 degradation is not necessary for AFP2 to repress ABA signalling in seeds. Surprisingly, *dwa1 dwa2* mutants overexpressing AFP2 displayed the unusual combination of desiccation tolerant viable seed with greatly reduced ABA sensitivity, indicating that these two characteristics can be molecularly separated. Using quantitative proteomics, we identified specific protein accumulation patterns in dry seeds correlating with one or both traits.

## Results

We initially tested for possible direct interactions between AFPs and E3 ligases by yeast two-hybrid and split YFP assays, including ABI5 as a positive control for interactions with both the DWA and KEG classes (Supplemental Figure S1). For the yeast two-hybrid assays, the E3 ligases were presented as GAL4 activation domain (AD)-fusions, and the AFPs and ABI5 were present in both AD- and binding domain (BD)-fusions. We focused on AFP1 and AFP2 because these had previously been shown to have the greatest effect on ABA sensitivity of germination (Lynch et al., 2017). As documented previously, the AFPs interacted with each other and ABI5 (Garcia et al., 2008), but both AD-DWA1 and AD-DWA2 fusions failed to interact with any BD-fusions. Although yeast expressing AD fusions to full-length KEG grew very poorly, even on non-selective media, AD fusions to the Ankyrin-HERC (AH) domains of KEG interacted with BD-fusions to ABI5 and the AFPs. The Ankyrin domains were previously shown to be essential for interactions with ABI5 (Stone et al., 2006). The split YFP assays also repeated strong interactions between ABI5 and the AFPs, but weaker interactions between DWA fusions and either ABI5 or the AFPs.

Although most interactions with the E3 ligases were weak and not consistent between these two assay systems, if functioning as part of a larger complex, these binary assay systems might not be sufficient to detect functional interactions. Furthermore, if the E3 ligase fusions encoded by these hybrid constructs are active, they may be marking any target fusions for degradation, resulting in reduced apparent interaction.

### Tests of epistasis between AFP overexpression and E3 ligase mutants

To test epistatic interactions, a *35S-YFP-AFP2* transgene was either transformed into the *dwa* mutants or crossed into the *keg* mutant backgrounds. Initial attempts to analyze interactions with *keg* knockout alleles were complicated by conflation of seed/seedling lethality that could be due to either loss of KEG or extreme AFP overexpression. Consequently, we crossed a previously characterized YFP-AFP2 overexpression line with an RNAi based knockdown line for *KEG* (Pauwels et al., 2015).

Analysis of multiple independent transgenic lines in the various *dwa* mutant backgrounds showed that AFP2 overexpression was fully capable of driving strong ABA resistance in this background, indicating that this effect was not dependent on DWA function (Figure 1A). Although the *KEG* knockdown did not completely prevent the AFP2 overexpression-induced germination on high ABA, germination was delayed by at least 1 day (Figure 1B).

**Figure 1.**
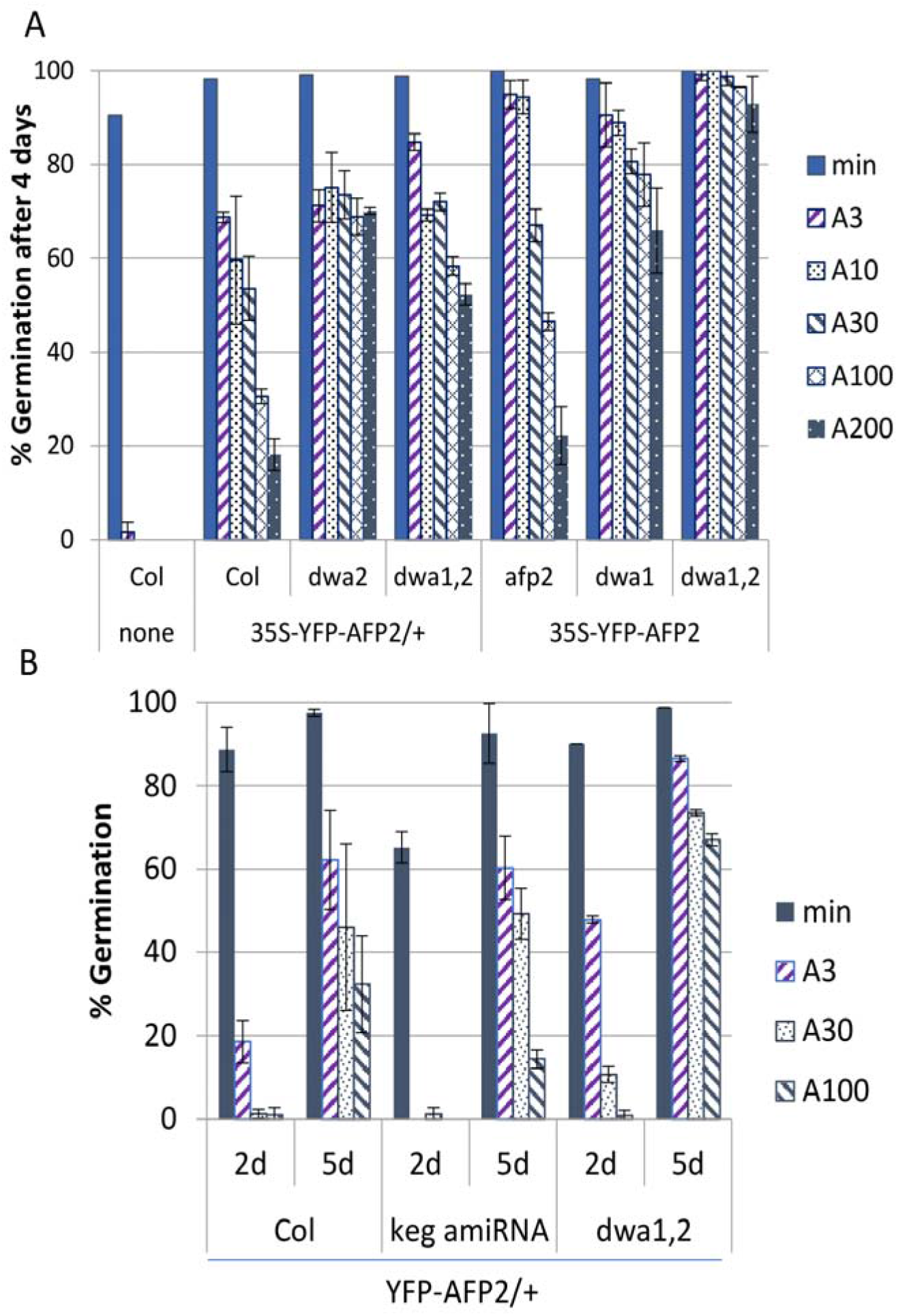
ABA sensitivity of germination for seeds overexpressing YFP-AFP2 fusions in wild-type (Col) or E3 ligase mutant (dwa) or knockdown (keg amiRNA) backgrounds. (A) Germination frequency after 4d post-stratification on media containing no ABA (min) or between 3 - 200 μM ABA (A3 to A200). (B) Germination frequency of hemizygous transgenic lines after 2-5d post-stratification on varying concentrations of ABA. Error bars = S.e.

To determine whether the observed ABA-resistant germination was due to either reduced ABI5 accumulation or extreme AFP overexpression, levels of these proteins were assayed by immunoblots of seed or seedling extracts over a time course of incubation on media with or without a concentration of ABA sufficient to inhibit germination of wild-type seeds (Figure2, Supplemental Figure S2). The YFP-AFP2 protein accumulated over the first 2d post-stratification in the wild-type and *dwa* mutant backgrounds (Figure 2A), roughly correlated with the extent of germination. In the *KEG* knockdown line, appearance of the YFP-AFP2 protein was slightly delayed, paralleling the delayed onset of germination (Figure 2B). Similar to previously documented changes in *ABI5* transcript level (Lynch et al., 2017), ABI5 protein levels decreased during stratification, then increased during continued exposure to ABA. As expected based on initial characterization of the *dwa* and *keg* mutants, ABI5 protein over-accumulated in the *dwa1 dwa2* mutant and *KEG* amiRNA backgrounds in the presence of ABA, correlating with inhibition of germination (Supplemental Figure S2). Surprisingly, in lines overexpressing YFP-AFP2, ABI5 was still present at high levels in seed populations that had reached up to 90% germination (Figure 2), indicating that the enhanced germination due to AFP2 overexpression was not due to loss of ABI5.

**Figure 2.**
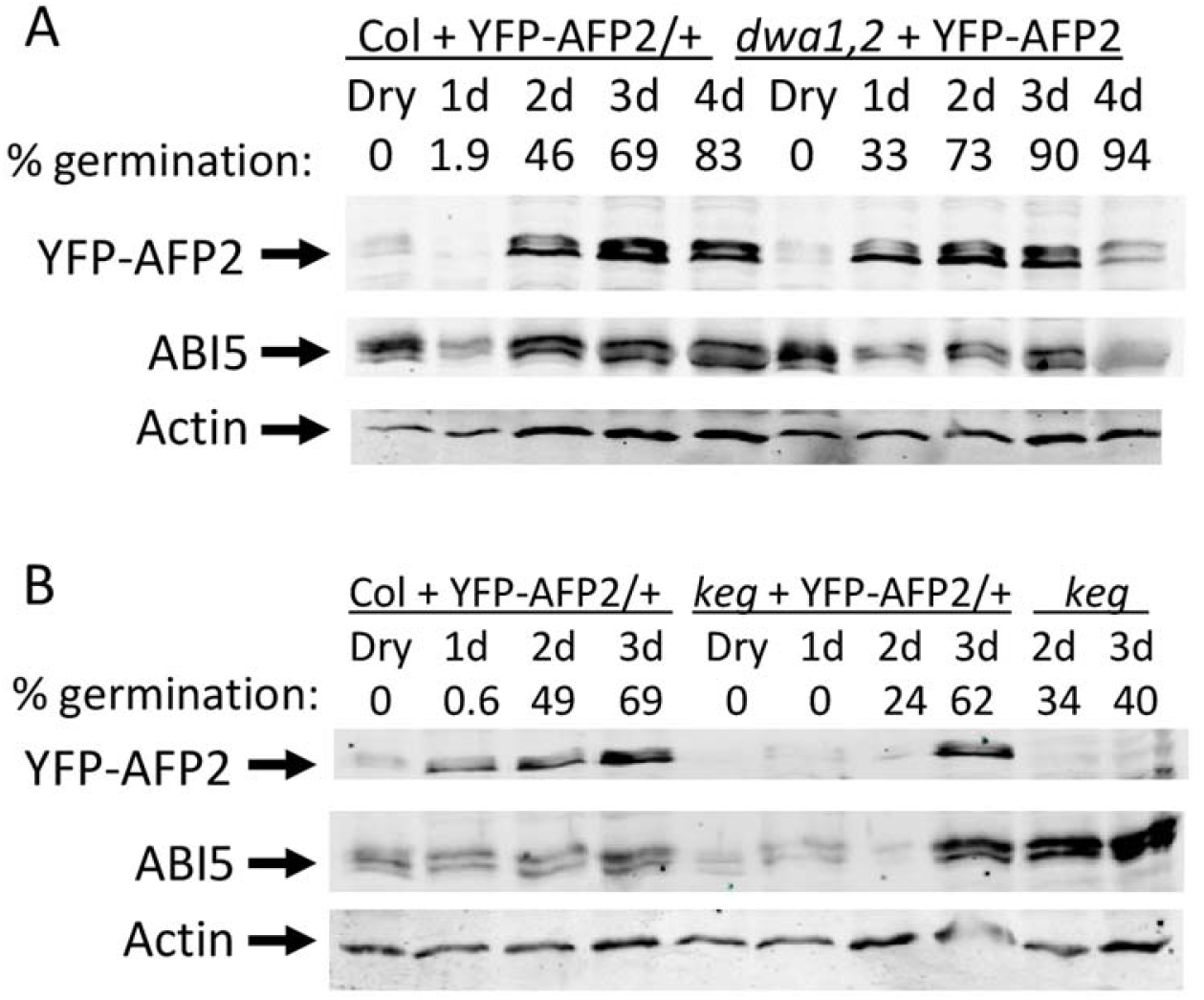
Timecourse of YFP-AFP2 and ABI5 accumulation and germination during incubation on GM + 1 μM ABA. Immunoblots were incubated with anti-GFP and anti-ABI5 antibodies to detect YFP-AFP2 and ABI5, respectively. Anti-actin was used as normalization control. (A) Comparison over 4d post-stratification in wild type (Col) vs *dwa1 dwa2* mutant backgrounds. (B) Comparison over 3d post-stratification in wild-type vs keg amiRNA backgrounds, and keg amiRNA parental line.

### Stability of ABI5 and AFP2 in wild-type or E3 ligase mutant backgrounds

We next sought to determine whether AFP2 overexpression could promote proteasomal degradation of ABI5 in seedlings deficient in the DWA or KEG E3 ligases. Transgenic lines with the YFP-AFP2 fusion in wild-type, *dwa1 dwa2* mutant or *KEG* knockdown backgrounds were incubated for 6d on media containing 1 μM ABA, a concentration low enough to maintain ABI5 accumulation without inhibiting germination. Seedlings were then incubated for an additional 6 hrs with or without cycloheximide or MG132, inhibitors of protein synthesis and proteasomal function, respectively. Consistent with published results (Lee et al., 2010; Stone et al., 2006), removal of ABA resulted in reduced ABI5 accumulation in all genotypes (Figure 3, lanes 1 vs. 2 for Col and *dwa* backgrounds, lanes 5 vs. 2 for *keg* background), but cycloheximide had this effect in only the wild type and *keg* knockdown lines (Figure 3, lanes 2 vs 3 for each genotype). This showed that, even when AFP2 was overexpressed, ABI5 was still stabilized in the *dwa1 dwa2* mutant background. Treatment with MG132 (lanes 4 vs 2 for each genotype) maintained ABI5 levels in the wild type and *keg* knockdown lines, and actually enhanced ABI5 accumulation in the *dwa1 dwa2* background. However, unlike the previously reported hyperaccumulation of ABI5 in *dwa1 dwa2* mutants incubated 5d on ABA (Lee et al., 2010), ABI5 levels were not higher when AFP2 was overexpressed in the *dwa1 dwa2* background than when overexpressed in the wild-type background. This suggested that ABI5 was still subject to some proteasomal degradation via a DWA-independent route.

**Figure 3.**
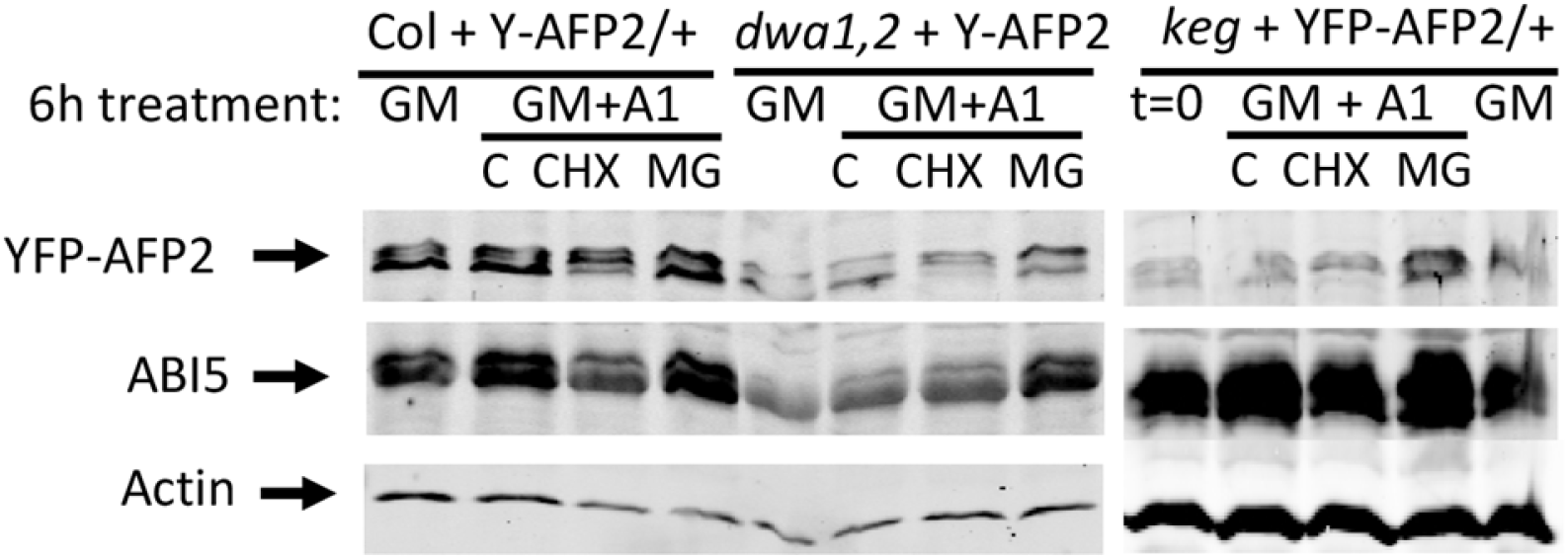
Stability of AFP2 in wild type vs. *dwa* or *keg* backgrounds. Seedlings were incubated 6 days post-stratification on GM + 1 μM ABA prior to 6h treatment in liquid GM, with or without 1 μM ABA. C=control, CHX = 100 μM cycloheximide, MG = 100 μM MG132. Anti-actin was used as a normalization control.

AFP2 itself is subject to proteasomal degradation when overexpressed as a YFP fusion (Figure 3), suggesting that it might be an additional substrate for one or more of these E3 ligases. We previously found that the YFP-AFP2 fusion migrates as a doublet on SDS-PAGE (Lynch et al., 2017), presumably due to post-translational modification. The highest mobility form of AFP2 is reduced in the presence of cycloheximide, indicating that it is less stable than a lower mobility form. Interestingly, treatment with cycloheximide results in accumulation of an intermediate mobility form, but this is blocked by treatment with MG132 which enhances accumulation of the lowest mobility form. Similar results are observed in the *dwa1 dwa2* mutant and *keg* amiRNA backgrounds, indicating that these E3 ligases are not required for destabilizing the high mobility form.

### Seed maturation and viability in Col vs *dwa* seeds overexpressing AFP2

Extreme ABA resistance in seeds is often correlated with failure to complete seed maturation, resulting in desiccation intolerant green seeds, as seen for null alleles of *ABI3* (Nambara et al., 1995) or lines strongly overexpressing AFP2 (Lynch et al., 2017). Consequently, lines overexpressing AFP2 in the Col-0 background were more readily maintained hemizygous for the transgene, and thus produced around 25% wild-type progeny. In contrast, overexpression in the *dwa1 dwa2* double mutant background did not disrupt acquisition of desiccation tolerance, so could be maintained as transgene homozygotes.

Chlorophyll fluorescence analysis of dry seeds overexpressing AFP2 showed an approximately 3-fold increase compared to Col-0 seed. In contrast, AFP2 overexpression in the *dwa* backgrounds, despite conferring strong resistance to ABA (Figure 1A), produced seeds with chlorophyll fluorescence like the Col-0 or the *dwa1 dwa2* parental line (Figure 4). Although transgenic lines in both Col-0 and the *dwa* double mutant backgrounds accumulated similar amounts of AFP2 fusion protein by 2 days after imbibition, the absence of DWA1 and DWA2 enhanced germination capacity from 25 to 68% on GM (Supplemental Figure S2).

**Figure 4.**
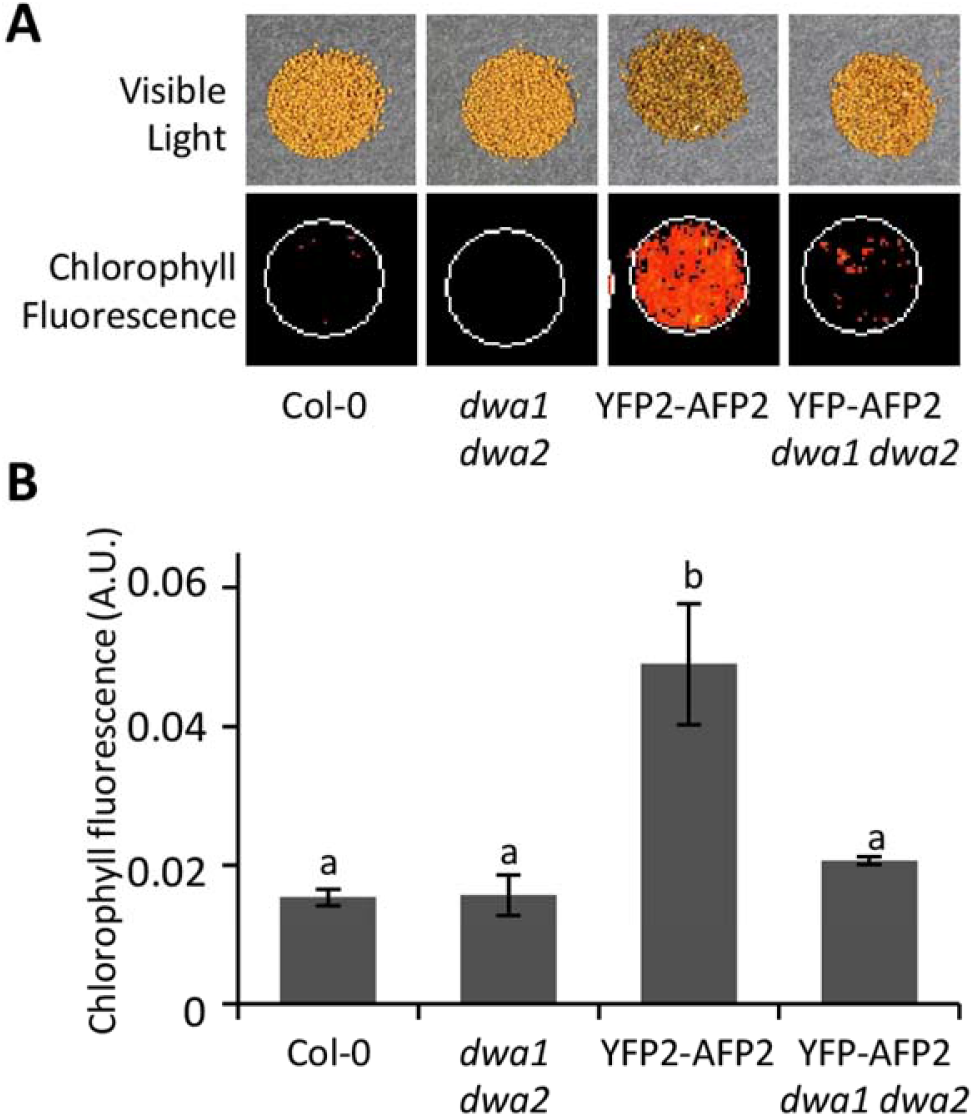
Chlorophyll accumulation in dry seeds. (A) Visible light (top) and Chlorophyll fluorescence (bottom) (B) Chlorophyll fluorescence quantification (One-way Anova-Tukey HSD test p=0.01 n=3).

### Proteomic analysis of AFP2 overexpression in Col vs *dwa* backgrounds

To identify changes in protein abundances correlated with improved viability of *AFP2* transgenic lines in the *dwa* compared to the Col-0 background, we performed a label-free quantitative mass spectrometry-based proteomic analysis comparing the dry seed of four genotypes in three independent biological replicates: Col-0, *dwa1 dwa2* double mutant, and YFP-AFP2 over expressing lines in both Col-0 (4A) and *dwa1 dwa2* (2A-2) backgrounds. In total 4070 protein groups were identified, including 3658 protein groups quantified in at least 2 replicates of a genotype (Supplemental Table S1). High reproducibility was observed across replicates (Supplemental Figure S3) and statistical analysis identified 1202 protein groups with significant (p<0.05, LIMMA statistics) abundance changes in at least one of the comparisons between genotypes (Supplemental Table S1). Of the reproducibly quantified proteins, only 2-11% were differentially accumulated in any given combination, with the greatest differences in proteins that were more abundant in the transgenic lines than the Col-0 and *dwa* backgrounds (Table 1).

**Table 1.** Extent of statistically significant changes in protein abundance in each pairwise comparison between genotypes.

| Genotype comparison | # proteins<br>(% of total<br>detected) | Enriched GO terms |
| --- | --- | --- |
| Col > *dwa1 dwa2* | 155 (4.2%) | Photosynthesis<br>Generation of precursor metabolites and energy<br>Response to abiotic, biotic and stress stimuli |
| Col > YFP-AFP2 | 183 (5%) | Embryonic development ending in seed<br>dormancy<br>Response to abiotic and stress stimuli |
| Col > YFP-AFP2, *dwa1 dwa2* | 109 (3%) | Response to abiotic and stress stimuli |
| *dwa1 dwa2* > YFP-AFP2,<br>*dwa1 dwa2* | 132 (3.7%) | Embryonic development ending in seed<br>dormancy<br>Response to abiotic and stress stimuli |
| YFP-AFP2 < YFP-AFP2, *dwa1*<br>*dwa2* | 176(4.8%) | Response to abiotic and stress stimuli<br>Secondary metabolic process<br>Response to biotic stimulus |
| Col < *dwa1 dwa2* | 112 (3.1%) | Translation |
| Col < YFP-AFP2 | 314 (8.6%) | Photosynthesis<br>Generation of precursor metabolites and energy<br>Lipid metabolic process<br>Cellular amino acid and derivative metabolic<br>process<br>Response to abiotic and biotic stimuli |
| Col < YFP-AFP2, *dwa1 dwa2* | 328 (9%) | Photosynthesis<br>Generation of precursor metabolites and energy<br>Response to cold |
| *dwa1 dwa2* < YFP-AFP2,<br>*dwa1 dwa2* | 393 (10.7%) | Photosynthesis<br>Generation of precursor metabolites and energy<br>Response to cold |
| YFP-AFP2 >YFP-AFP2, *dwa1*<br>*dwa2* | 82 (2.2%) | Lipid metabolic process |

We performed a clustering analysis using the 1202 significantly regulated proteins and identified six clusters that correlated with differences in ABA sensitivity or desiccation tolerance (Figure 5A; Supplemental Table S2). Clusters I and VI corresponded to proteins down- and up-regulated, respectively, only in green seeds, maintaining the usual correlation between extreme ABA resistance and poor viability. Clusters II and V included proteins decreased or increased, respectively, in all ABA-resistant seeds, independently of maturation defect phenotypes. Clusters III and IV represented protein changes unique to seeds displaying the unusual combination of ABA resistance and good viability. We performed GO analyses with proteins from each subgroup (Figure 5B; Table S3). Cluster I (n=50) was enriched in regulation of GTPases, whereas cluster VI (n=43) was enriched in the categories of fatty acid biosynthesis and nuclear import. Cluster II (n=137) and V (n=356) were both enriched in temperature related stress but in distinct heat or cold subcategories. While Cluster II was the only sub-group with enrichment in response to ROS and to a lesser extent in seed maturation, cluster V was additionally enriched in osmotic, salinity, and pathogen stress responses but especially in GO terms associated with growth: photosynthesis, generation of precursor metabolites (amino acids, nucleotides, lipids and carbohydrates) and energy. Overaccumulation of these proteins reflects the failure to arrest primary metabolism during maturation, hence increasing their germination potency but not supporting extended viability. Cluster III (n=27) did not lead to significant enrichment due to its limited size but included proteins related to defense response, lipid metabolism/storage, and LEAs. Cluster IV (n=79) was enriched mostly in cold-associated stress response proteins.

**Figure 5.**
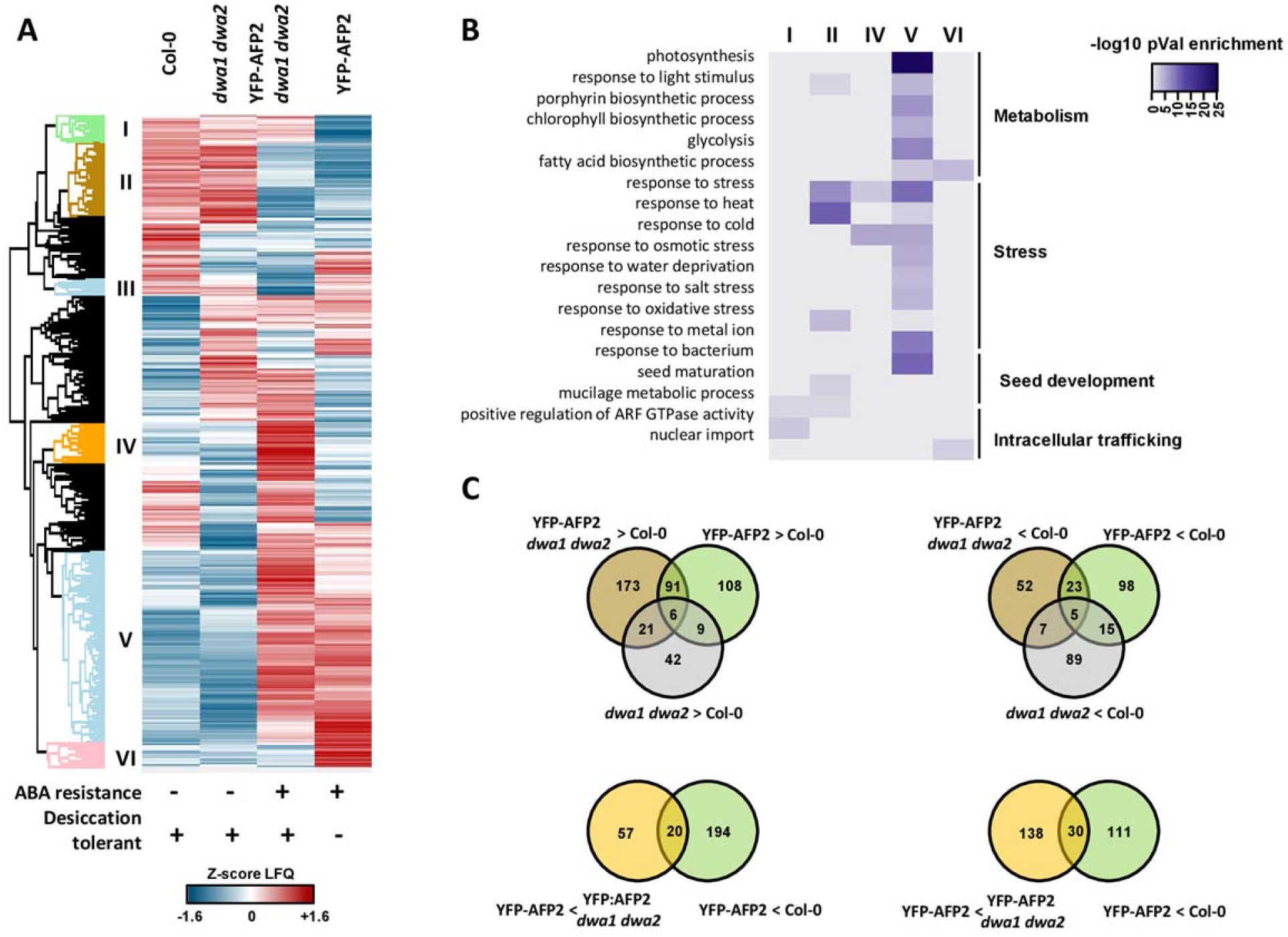
Proteome comparisons highlighting overlap correlated with ABA resistance. (A) Heat map of Z scores for all proteins identified as significantly differentially accumulated in at least one comparison(B) Enrichment of GO categories in protein clusters identified in heat map. (C) Venn diagrams showing overlaps between proteins over-accumulated or under-accumulated in *dwa1,dwa2* and/or YFP-AFP2 overexpression lines relative to wild-type (Col-0) or between YFP-AFP2 overexpression in Col-0 vs *dwa1,dwa2* backgrounds.

To identify changes of high amplitude associated with ABA resistance and/or desiccation tolerance, we searched for overlaps between genotypes of protein significantly regulated by at least a two-fold change in comparison to Col-0 (Figure 5C; Supplemental Table S4). We found 91 proteins increased, and 23 proteins decreased, in seeds overexpressing YFP-AFP2 in either background, but not differentially accumulated in *dwa1 dwa2 vs*. Col-0 backgrounds. Differential accumulation of these proteins correlated with ABA resistance, and included increases in proteins associated with photosynthesis and growth, and decreases in some seed-specific proteins. Comparisons of proteins differentially accumulated between seeds overexpressing YFP-AFP2 in *dwa1 dwa2* vs. Col-0 backgrounds identified 108 proteins that were more abundant, and 98 that were less abundant exclusively in the green seed batches; increases were again enriched for photosynthetic functions (partially overlapping clusters V and VI), while decreases were enriched for storage reserve proteins (partially overlapping clusters I and II). Differential accumulation specific to seeds with the unusual combination of desiccation tolerance and extreme ABA resistance during germination included 173 and 52 proteins that were up- and down-regulated, respectively; increases were enriched for response to cold, while decreases included some heat shock proteins (HSPs) and ABI5. Further comparisons of proteins differentially accumulated in YFP-AFP2 overexpressors relative to either Col-0 or YFP-AFP2 overexpression in the *dwa1 dwa2* background identified 20 proteins that were decreased, and 30 proteins that increased, in the desiccation tolerant seeds (Figure 5C).

The accumulation of some highly abundant seed specific proteins during maturation was affected by overexpression of AFP2 but not all members of any given class of proteins were similarly affected. For example, although 29 Late Embryogenesis Abundant (LEA) proteins were detected in the analysis, only 9 showed significantly reduced accumulation in at least one of the YFP-AFP2 overexpression lines compared to the wild type (Supplemental Figures S4A,B). When comparing the effect of the AFP2 transgene in the *dwa* background against its parental line, 10 LEAs were found to be significantly reduced, but two plastid-localized LEAs were significantly increased in this comparison (Supplemental Figure S4D). All three Cruciferins, the major type of seed storage protein in Arabidopsis, were down regulated by AFP2 over expression together with At2S3 but this regulation was abolished by the *dwa* mutations (Supplemental Figures S4A,B). Three oleosins showed significant regulation by AFP2 overexpression, but the pattern was inconsistent across comparisons (Supplemental Figure S4). This illustrates the complexity by which overexpression of AFP2 affects seed quality.

In summary, this analysis identified several subsets of proteins for which differential accumulation is associated with extreme resistance to ABA during germination and/or seed viability including component(s) putatively directly involved in the functions of AFP2 and/or DWAs regarding the control of seed quality.

## Discussion

One or more of the AFPs have been identified in multiple screens for Arabidopsis proteins interacting with ABI5 (Lopez-Molina et al., 2003; Garcia et al., 2008; Lumba et al., 2014; Chang et al., 2018) or their homologs in other species (Tang et al., 2016). Although highly conserved across multiple domains and showing some redundant functions, they are not fully coordinately regulated and also have some distinct functions. Three of the four Arabidopsis AFPs (AFP1, AFP2 and AFP3) are negative regulators of ABA response (Garcia et al., 2008) and all are induced by ABA or a variety of stress treatments.

### AFP expression, interactions, roles and proposed mechanisms

All of the AFPs are expressed in seeds and seedlings, with AFP2 expressed at higher levels in maturing and dry seeds, and AFP1 and AFP3 becoming more highly expressed in response to stresses during vegetative growth. Consistent with these expression patterns, genetic analyses have shown that AFP1 and AFP2 act redundantly to reduce ABA sensitivity at germination (Garcia et al., 2008) and AFP2 breaks high temperature-induced secondary dormancy via effects on GA and ABA metabolism by suppressing ABI5- and DELLA-mediated SOMNUS (SOM) expression (Chang et al., 2018). In addition, AFP2 attenuation of responses to salt and osmotic stress may be mediated via interactions with SnRK1 kinases (Carianopol et al., 2020), and AFP2 can promote cell death when transiently overexpressed (Carianopol and Gazzarrini, 2020). Later in development AFP2 and AFP3 act redundantly to inhibit flowering under long day conditions by reducing expression of *CO* and its downstream targets *FT* and *SOC1* (Chang et al., 2019).

AFP1 was initially proposed to attenuate ABA and stress responses by acting as an adaptor for E3 ubiquitin ligases that target ABI5 and related bZIPs for destruction via the proteasome (Lopez-Molina et al., 2003) and this model has been extended to apply to all of the AFPs (Zhang et al., 2019). However, analyses of *afp2* mutants showed that ABI5 protein levels in the mutants were higher than those in the wild-type seeds only under conditions when the wild-type seeds had germinated but the mutants had not, suggesting that ABI5 degradation was an effect rather than a cause of germination (Garcia et al., 2008). Subsequent studies showed that these AFPs interact with TOPLESS (TPL), some TOPLESS-RELATED proteins (TPRs), and histone deacetylase (HDAC) subunits, consistent with modulating ABA response in part via effects on chromatin modification, similar to the mechanism of jasmonate signaling repression by NINJA (Pauwels et al., 2010; Lynch et al., 2017). With respect to control of flowering, AFP2 interacted directly with both CO and TPL or TPR2 to form a complex that recruited HDAC activity to reduce acetylation of the FT promoter, thereby reducing FT expression and delaying flowering (Chang et al., 2019). AFP2 effects on flowering time also appeared to be correlated with proteasomal degradation of CO, consistent with multiple mechanisms of action. Similar interactions in rice between MODD and OsbZIP46, homologs of AFP and ABI5 respectively, resulted in decreased expression of ABA- and drought-induced gene expression *via* reduction of both OsbZIP46 stability and histone acetylation of OsbZIP46 target genes (Tang et al., 2016).

To test whether overexpressed AFP2 was promoting germination by enhancing proteasomal degradation of ABI5, we assayed ABI5 accumulation in wild-type and E3 ligase mutant backgrounds. Previous studies have shown that ABI5 accumulates during seed development, decreases transiently during stratification, and either continues to decline during germination or increases back to high levels during exposure to ABA or stress treatments (Lopez-Molina et al., 2001). Seeds with mutations in KEG or the DWAs, E3 ligases shown to ubiquitinate ABI5, maintain ABI5 at high levels even under low or no ABA or stress conditions that are not sufficient to block germination of wild-type seeds (Stone et al., 2006; Lee et al., 2010). This is correlated with the mutants’ failure to germinate, but does not distinguish between maintenance of ABI5 as a cause or effect of non-germination. In the current work, YFP-AFP2 fusion expression was controlled by the CaMV 35S promoter which, although considered constitutive, is expressed more highly post-germination than in seeds of several species (Terada and Shimamoto, 1990). Consequently, increased YFP-AFP2 accumulation accompanied germination. However, the YFP-AFP2 level present at seed maturity was very much higher than the respective background levels for the endogenous AFP2 protein in *dwa1 dwa2* and Col seeds (Supplementary Table S1) and was sufficient to dramatically increase germination despite maintaining ABI5 protein at high levels until germination was nearly complete, regardless of whether KEG or the DWAs were functional. This shows that loss of ABI5 coincides with germination, but does not need to precede it. Furthermore, the germination promoting activity of AFP2 did not depend on activity of these E3 ligases, so it does not appear to act as an adaptor for them, at least under these conditions. Although it is possible that the loss of one class of E3 ligase may be compensated by the continued activity of the other class, the strong ABA-hypersensitive phenotypes of the *dwa* and *keg* mutants lacking AFP2 overexpression argue against this.

Yeast two-hybrid and BiFC tests of direct interactions between the AFPs and E3 ligases were ambiguous, showing at most weak interactions in either assay. However, even ABI5 appeared to interact weakly with these E3 ligases in these assays, and published studies have demonstrated that ABI5 is a direct substrate for both the DWA and KEG class E3 ligases (Lee et al., 2010; Stone et al., 2006). Immunoblot analyses, combined or not with the use of cycloheximide (CHX), showed that the high mobility form of AFP2 was proteasomally degraded in all three genetic backgrounds, suggesting that it is a substrate for a different class of E3 ligases.

### Seed omics

Comparisons of seed transcriptome and proteome data have shown extensive post-transcriptional regulation and post-translational modifications (Mergner et al., 2020). In seeds selective translation of mRNA and post-translational modifications play prominent roles in the control of germination *per se* (Arc et al., 2011; Sano et al., 2020; Galland and Rajjou, 2015). Therefore, the dry seed proteome provides a comprehensive view of the final products of gene expression, which define seed traits. Although desiccation tolerance is necessary for longevity of orthodox seeds, these traits may be acquired successively during seed maturation and regulated differently (Buitink and Leprince, 2018). Desiccation tolerance, whether in seeds or vegetative tissues, requires protection from protein denaturation, changes in membrane conformation, and oxidative damage to lipids and proteins (Oliver et al., 2020). These are associated with accumulation of compatible solutes, including non-reducing sugars, LEA proteins, dehydrins and HSPs, and increases in antioxidant enzymes. Seed longevity is further correlated with glass formation, accumulation of molecular antioxidants, loss of chlorophyll, and production of proteins needed to prevent damage to DNA and RNA. Omics approaches have identified defense-related compounds associated with longevity, possibly as a mechanism for pre-emptive protection from pathogen attack in a dormant or quiescent seed.

Within the numerous available seed proteomic reports (Narula et al., 2016), studies directly aiming at the identification of protein involved in seed longevity/viability present in dry seeds unveiled 309 differentially accumulated proteins in GERMINATION ABILITY AFTER STORAGE (GAAS) lines (Nguyen et al., 2015), 320 in *mdn1-1* (Li et al., 2019), and 442 in *vps29* mutants (Durand et al., 2019). The study using GAAS lines exploited natural variation in seed storability and proposed that the highly abundant Cruciferin storage proteins (CRA, CRB and CRC) act as a ROS buffering system (Nguyen et al., 2015). MIDASIN 1 (MDN1) is involved in ribosome biogenesis and, although a null allele is embryo lethal, the *mdn1-1 (dsr1)* weak allele mutant produces large seeds with low germination rates that can be partially rescued by loss of ABI5 function, compensating for the overexpression of ABI5 in this mutant. Vacuolar Protein Sorting (VPS) 29 is a subunit of the retromer, involved in protein trafficking and recycling within cells; a null mutant has greatly reduced seed vigor and longevity, but does not affect desiccation tolerance. Nearly 90% of the proteins identified in the study of *vps29* were also detected in the current study; approximately one-third of these were differentially accumulated. Changes that were correlated with reduced seed vigor included decreases in oxidation-reduction processes and response to light and temperature, but increases in translation, vesicle-mediated transport, fatty acid metabolism, proteolysis, and response to abiotic stresses and ABA. Using mutants affected in their seed ABA content by impairment of biosynthesis or catabolism during maturation, (Chauffour et al., 2019) identified 120 proteins with changed accumulation. Approximately 40% of these proteins were found differentially accumulated in the current study. Consistent with previous studies of seed ABA response, these included ABA-dependent accumulation of many proteins associated with storage reserves and desiccation tolerance, and reduction of those involved in reserve mobilization, photosynthesis, cell cycle, and protein oxidation.

ABA signaling promotes acquisition of desiccation tolerance and dormancy during seed maturation, limits germination to environmental conditions that favor seedling establishment, and is important for imposing an ABI5-mediated developmental checkpoint preventing post-germination growth under adverse conditions (reviewed in (Finkelstein, 2013; Ali et al., 2021). In our study, we took advantage of the extreme ABA resistance of AFP2 overexpressing seeds, and their restored viability in the context of *dwa1 dwa2* mutations, to identify 1202 significant changes of protein abundance in dry seed across pairwise comparisons of four genotypes. We found 6 protein accumulation patterns and several sets of strongly regulated proteins correlating with 2 seed traits (extreme ABA resistance and green seed phenotype) in combination or independently. Analysis of regulated biological processes reinforces and refines the classes described in the earlier studies. ABA resistance correlates with increased production of proteins associated with photosynthesis and growth (Cluster V) and decreases in those associated with stress response proteins such as HSPs, LEAs, defense proteins, and storage proteins implicated in protection from oxidative damage (Cluster II). This last cluster also included an osmotic stress activated kinase, SnRK2.10, and the transcription factor SOMNUS, which is activated by ABI3 and ABI5 and in turn indirectly promotes ABA accumulation (Supplemental Figures S4 C, E) (reviewed in (Sano and Marion-Poll, 2021).

Our analysis further indicates a specific role of cellular trafficking events in the control of seed desiccation tolerance independently of ABA resistance (cluster I), with a possible role of NEVERSHED (NEV)(down-regulated in a desiccation sensitive genotype), an ARF GTPase controlling floral abscission (Stefano et al., 2010; Groner et al., 2016). Although no seed phenotype has been reported yet for *nev* mutants, likely due to redundancy within the family, other mutants affected in vesicle transport display reduced seed longevity (Zhao et al., 2018; Durand et al., 2019). Conversely, repression of nuclear import is associated with better seed storability (Cluster VI), in that components of the protein nuclear import complex, SIRANBP and NUCLEAR TRANSPORT FACTOR2 (NTF2) (Meier and Brkljacic, 2010), are over-accumulated in green seeds (Supplemental Figure S4A). The current study identified additional proteins for which low accumulation correlates with a desiccation intolerant state (Cluster I) that had not been found in the earlier proteomic studies. These included the photoreceptor PHYTOCHROME E, previously shown to contribute to R/FR regulation of germination (Hennig et al., 2002) and YODA, a MAP3K regulating seed development (Lukowitz et al., 2004). Further changes included increases in regulators such as the chromatin remodeling protein SWI3D (Sarnowski et al., 2005) and the wall remodeling protein XYLOGLUCAN ENDOTRANSGLUCOSYLASE/HYDROLASE 19. In contrast, TARGET OF RAPAMYCIN (TOR), which promotes germination and growth in part through antagonisms with ABA signaling (Fu et al., 2020), was reduced in the *dwa* background, whether or not the *YFP-AFP2* transgene was present.

Proteins over-accumulated exclusively in the unusual combination of extreme ABA resistance and seed storability (Cluster IV) include a GIBBERELLIN METHYLTRANSFERASE, which is normally expressed during seed maturation, leading to inactivation of GA (Varbanova et al., 2007), MIDASIN1 (MDN1) previously shown to be required for a proper shaping of the seed maturation proteome (Li et al., 2019), and a Pumilio RNA binding protein (APUM24), for which loss of function leads to aborted seeds (Maekawa et al., 2017). These seeds also over-accumulate a beta-glucoside that releases ABA from a conjugated form (AtBG1) (Lee et al., 2006) and is up-regulated by ABA-deficiency (Chauffour et al., 2019). Failure to either produce or respond to ABA often has similar effects. Proteins with reduced accumulation in this combination of traits (Cluster III) include CORONATINE INSENSITIVE SUPPRESSOR1 (COS1) (Xiao et al., 2004) suggesting that positive regulation of JA signaling during maturation is associated with low seed storability.

Finally, to explore the molecular basis of the restored viability when YFP-AFP2 is overexpressed in the *dwa1 dwa2* background, we identified sets of proteins for which the desiccation tolerant lines, Col-0 and *YFP2-AFP2 dwa1 dwa2*, show similar differences relative to AFP2 overexpression in the Col-0 background (Fig. 5C). Within the 20 proteins under-accumulated, relative to YFP-AFP2 overexpression in the wild-type background, was a member of the CROWDED NUCLEI (CRWN) family that promotes ABI5 degradation (Zhao et al., 2016) as well as the Allene Oxide Synthase (AOS) implicated in JA biosynthesis, again consistent with maintaining ABA signaling and inhibiting JA-related signaling in desiccation tolerant seeds. Lastly within the 30 proteins up regulated, we identified a Pumilio RNA binding protein (APUM5), predicted to be localized in the nucleus, that is highly expressed in seeds and is associated with both biotic and abiotic stress responses (Huh and Paek, 2014). Interestingly, PUM5 has been shown to be ubiquitinated at position Lys55 suggesting that its stoichiometry is at least partly controlled by E3 ligase(s) and the 26S proteasome (Walton et al., 2016).

In summary, our study shows that although AFP2 can interact with E3 ligases that target ABI5 for proteasomal destruction, neither those E3 ligases nor ABI5 destruction is required for AFP2 overexpression to promote germination. This may reflect the very high ratio of AFP2:ABI5 in the overexpression lines, a quantity previously suggested to act as a “sensor” controlling germination (Garcia et al., 2008). The slight delay in ABA-resistant germination observed in the 35S-YFP-AFP2 *keg* amiRNA seeds is consistent with the delay in AFP2 accumulation in this line. Finally, the proteome comparison identified subsets of proteins that correlated with good desiccation tolerance and viability despite greatly reduced ABA sensitivity. Although superficially contradictory, these combinations of altered protein expression reflect the balance between a predisposition to germinate and a stable ungerminated state, physiologically similar to seed “priming.” This information may be used to further dissect molecular mechanisms of desiccation tolerance and/or acquisition of ABA resistance during maturation, or as a reference for characterizing seed batch quality.

## Materials and Methods

### Plant Materials and Transgenes

Arabidopsis plants were grown in pots in growth chambers under continuous light at 22°C. *DWA* and *KEG* loss of function lines (*dwa1-1, dwa2-1, dwa1-1 dwa2-2, keg-1*, and *KEG* amiRNA) were described in (Lee et al., 2010; Stone et al., 2006; Pauwels et al., 2015). The *35S:YFP:AFP2* fusion and split YFP fusions for ABI5, AFP1 and AFP2 were described in (Lynch et al., 2017). *Agrobacterium tumefaciens*-mediated direct transformation of *dwa* mutants was performed by the floral dip method (Clough and Bent, 1998), followed by selection of BASTA-resistant seedlings. Homozygous lines were identified by production of 100% BASTA-resistant progeny. Following a cross between a *35S:YFP:AFP2* fusion line and *KEG* amiRNA line #14, YFP-AFP2 overexpressing progeny were selected by BASTA resistance and homozygous *KEG* knockdown segregants were identified by red fluorescence of seeds due to *ProOLE1:OLE1-RFP* expression.

Split YFP fusions for DWA1 and DWA2 were constructed using the Gateway compatible pSITE-nEYFP-C1 (GenBank Acc# GU734651) and pSITE-cEYFP-C1 (Acc# GU734652) vectors and PCR products with attL ends added as described in (Fu et al., 2008), following manufacturer’s instructions for LR Clonase reactions (Invitrogen). BiFC assays were conducted as described in (Lynch et al., 2017).

### Yeast Two-Hybrid Constructs and Assays

Fusions between the GAL4 activation domain (AD) and full-length DWA cDNAs were constructed using the pGADT7-DEST vector (Lu et al., 2010) and PCR products with attL ends. Fusions with full-length KEG and subdomains of KEG were constructed by LR Clonase reactions with the pGADT7-DEST vector and pDONR clones.

### Plant growth conditions

Germination assays testing ABA sensitivity of age-matched seeds were performed on minimal nutrient media supplemented with ABA at concentrations over the range from 0-200 μM, as described in (Lynch et al., 2017). Accumulation of fusion proteins was assayed by immunoblots of seeds or seedlings harvested after 0-4d incubation on Germination Medium (GM: 0.5x MS salts and vitamins, 1%sucrose) supplemented with 1 μM ABA and solidified with 0.7% agar. For testing stability of fusion proteins, seedlings were grown initially on solid GM containing 1 μM ABA, then transferred to liquid GM in multi-well plates with or without ABA, cycloheximide, MG-132 (Peptides International) or the appropriate solvent controls at the concentrations indicated.

### Immunoblots

Seeds or seedlings were ground directly in 1x or 2x Laemmli loading buffer, respectively, microfuged 10 min at 4°C to pellet debris, then boiled 5 min prior to fractionation by SDS-PAGE. Proteins were transferred to nitrocellulose filters, as described in (Lynch et al., 2017). Filters were blocked with Casein blocking buffer (LI-COR Biosciences, Lincoln, NE), then co-incubated with anti-GFP mAb(1:10000, UBPBio, Aurora, CO) and anti-ABI5pAb (1:10000, Ab98831, AbCam) primary antibodies, followed by anti-mouse and anti-rabbit secondary IRDye 800 conjugated IgGs, and visualized using the 800 channel of theLicor Odyssey Infrared Imaging System. Filters were subsequently probed with anti-actin mAb (A0480, Sigma), followed by anti-mouse secondary IRDye 800 conjugated IgGs.

### Dry seed proteome analysis

Approximately 25 mg of dry seeds (12 days after harvesting) per biological replicate and genotype were frozen in liquid nitrogen and ground to a fine powder using mortar and pestle. Total protein was extracted in 50 mM Hepes (pH 7.5), 3% SDS, 10 mM DTT, and 1:100 v/v protease inhibitor cocktail (P9599 Sigma-Aldrich) under agitation for 20 min. 12 ug of protein were diluted in 8 M urea and 0.1 m Tris/HCl, pH 8.5 and reduced by adding 10 mM DTT for 1 hour (Xiang et al., 2016). Reduced thiols were alkylated by addition of chloroacetamide (CAA) 45 mM final concentration for 1 hour under agitation in the dark. Excess of CAA was quenched by addition of DTT (83 mM final concentration and incubation for 1 hour). Sample was diluted 7 times with 50 mM ammonium bicarbonate buffer to reduce urea concentration. Trypsin (in solution) digestion (1:100 enzyme/protein) was performed over night at 37°C. Digestion was stopped by acidification with trifluoroacetic acid (TFA) (1% final concentration), the resulting mixture was loaded on centrifugation units (Amicon Ultracel-10, Millipore). Peptides were recovered in flow-through after centrifugation. The membranes were washed with 0.5 M NaCl and flow-throughs were combined. Peptides were desalted and pre-fractionated (in 3 fractions) by stage-tipping using Empore Styrenedivenylbenzene Reversed Phase Sulfonate material (SDB-RPS; 3M) (Kulak et al., 2014). Peptides were then dried and resuspended in 2% ACN, 0.1% TFA before MS measurement. Samples were analyzed on an EASY-nLC 1200 system (Thermo Fisher Scientific) coupled to a Q Exactive HF Orbitrap mass spectrometer (Thermo Fisher Scientific). Mass spectra were acquired in data-dependent acquisition mode with a TOP15 method as previously described (Bienvenut et al., 2020). MS/MS spectra were searched using MaxQuant software (v. 1.6.17.0) (http://www.maxquant.org/) against the Arabidopsis Araport11 database (Krishnakumar et al., 2014) with label-free quantification (LFQ) enabled (Cox et al., 2014) with standard settings. Maxquant output files were proceeds using Perseus (V1.6.6.0) (Tyanova et al., 2016). LFQ intensities were log2 transformed and data filtered for valid values in at least 2 out the 3 replicates of one genotype. Missing values were imputed from normal distribution using a width of (log2) 0.55 and a downshift of (log2) 1.9 (Supplementary Figure S6). Significant changes in protein abundances were calculated using the LIMMA (“Linear Models for Microarray Data”) package on R (Kammers et al., 2015). Each comparison set was further filtered to identify proteins with p-value ≤0.05 between any given pair of genotypes. Heat maps were built using Z-scored LFQ values in Perseus. Overlaps of significantly regulated proteins were obtained using Venny (Oliveros, 2007-2015), and indicated subsets were analyzed by Cytoscape, AgriGOv2.0 (Tian et al., 2017), and manually by identification of predicted products (Reiser et al., 2017).

## Data availability

The mass spectrometry proteomics data have been deposited to the ProteomeXchange Consortium (http://proteomecentral.proteomexchange.org) via the PRIDE partner repository with the data set identifier PXDxxxxx.

## Acknowledgements

We thank Dr. Xing-Wang Deng for the *dwa* mutant lines and DWA cDNAs, and Dr. Judy Callis for the *keg* mutant and amiRNA lines and KEG fusion and cDNA clones. In addition, we thank Paulina Heinkow and Dr. Jürgen Eirich for support with the MS analyses. pGADT7-GW was a gift from Yuhai Cui (now available as Addgene plasmid # 61702; http://n2t.net/addgene:61702; RRID:Addgene_61702). This work was supported by UCSB Academic Senate Grant to RRF, Faculty Research Assistance Program funds, National Science Foundation Grant# IOS1558011 to RRF, the Deutsche Forschungsgemeinschaft Grant# NE2296 to GN and INST211/744-1 to IF. TK was supported by a fellowship of the Studienstiftung des deutschen Volkes.

## Supplementary Figures

Fig. S1 Yeast two-hybrid and bimolecular fluorescence complementationassays of interactions between AFPs and DWA or KEG. Haploid yeast carrying plasmids encoding the indicated GAL4 AD or BD (pGBKT7) fusions were mated on YPD, then replica plated to plates selecting for either both plasmids (-LW) or interactions promoting reporter expression (-HALW). Fusions to the N- and C-terminal halves of YFP were co-infiltrated into *N. benthamiana* leaves. Micrographs of the lower epidermis were taken 2-3 days after infiltration.

Fig. S2 Western blots comparing ABI5 and YFP-AFP2 accumulation in parental vs. transgenic lines at 1 or 2 days post-stratification on GM with or without 1 μM ABA. Actin was used as a normalization control.

Fig. S3 Scatter plots for quality analysis of the seed proteome replicates. Log2-transformed Label Free Quantitative (LFQ) intensity values from two replicates are plotted against each other. Pearson correlation coefficients (Pcc) are shown on each panel.

Fig. S4 Volcano plots from quantitative proteomic analysis. Each dot corresponds to a unique protein group. The log2-fold change of protein abundances between genotypes are plotted against the p-value (-log10). Blue and red dots indicate proteins with significant down- and up-regulation, respectively. Proteins discussed in the text are highlighted in orange and labelled. The naming of LEA and Oleosins proteins follow the nomenclature from (Candat et al., 2014; Kim et al., 2002).

Fig. S5 Distribution of log2 LFQ values intensities before (A) and after (B) imputations. Imputations were performed from normal distributions of each samples with a downshift and a width of 1.9 and 0.55, respectively. Imputed LFQ intensities are highlighted in red.

## Supplementary Tables

Supplementary Table 1. List of 3658 quantified protein groups quantified in at least two biological replicates of a genotype and LIMMA statistical analyses.

Supplementary Table 2. List of proteins from the six clusters correlating with ABA resistance and/or seed viability.

Supplementary Table 3. Significantly enriched GO terms and lists of associated proteins for the six clusters correlating with ABA resistance and/or seed viability.

Supplementary Table 4. Overlap between genotypes of proteins significantly regulated by at least a twofold change.

## Notes

**Funding information:** This work was supported by UCSB Academic Senate Grant to RRF, Faculty Research Assistance Program funds, National Science Foundation Grant# IOS1558011 to RRF, the Deutsche Forschungsgemeinschaft Grant# NE2296 to GN and INST211/744-1 to IF. TK was supported by a fellowship of the Studienstiftung des deutschen Volkes.

